# Express saccades during a countermanding task

**DOI:** 10.1101/2020.03.20.000760

**Authors:** Steven P. Errington, Jeffrey D. Schall

## Abstract

Express saccades are unusually short latency, visually guided saccadic eye movements. They are most commonly observed when the fixation spot disappears at a consistent, short interval before a target spot appears at a repeated location. The saccade countermanding task includes no fixation-target gap, variable target presentation times, and the requirement to withhold saccades on some trials. These testing conditions should discourage production of express saccades. However, two macaque monkeys performing the saccade countermanding task produced consistent, multimodal distributions of saccadic latencies. These distributions consisted of a longer mode extending from 200 ms to as much as 600 ms after target presentation and another consistently less than 100 ms after target presentation. Simulations revealed that by varying express saccade production, monkeys could earn more reward. If express saccades were not rewarded, they were rarely produced. The distinct mechanisms producing express and longer saccade latencies were revealed further by the influence of regularities in the duration of the fixation interval preceding target presentation on saccade latency. Temporal expectancy systematically affected the latencies of regular but not of express saccades. This study highlights that cognitive control can integrate information across trials and strategically elicit intermittent very short latency saccades to acquire more reward.

## INTRODUCTION

Saccade latency is a manifestation of visual, motor, and cognitive processes (Carpenter 1988). Saccade latency and dynamics are influenced profoundly by reward expectancy and value. Saccades to stimuli associated with higher reward typically are more accurate and have faster peak velocities and shorter latencies relative to unrewarded stimuli (Milstein and Dorris 2007; Takikawa et al. 2002; Vullings and Madelain 2018; 2019). Of course, reward contingencies are integral in learning behaviors, and previous work has highlighted that production of express saccades is learnt over time (Bibi and Edelman 2009; Fischer et al. 1984; Johannesson et al. 2018; McPeek and Schiller 1994; Pare and Munoz 1996).

Saccade latency is also influenced by the reliability of the timing of events. Behavioral testing typically includes an interval between the presentation of a warning stimulus and the presentation of a target stimulus, known as the foreperiod. The passage of time within a foreperiod can convey information about when to expect the target. Sampling foreperiods from a uniform rectangular distribution results in an aging distribution with the conditional probability of the target appearing increasing as the foreperiod elapses. Conversely, non-aging foreperiods have an exponentially declining probability of elapsing, resulting in a constant conditional probability. Response times, including saccade latencies, are typically quicker following predictable, longer foreperiods (Ameqrane et al. 2014; Correa and Nobre 2008; Drazin 1961; Naatanen 1970; Niemi and Näätänen 1981; Thomaschke et al. 2011). Saccade latencies can become extremely short following a brief (~200 ms) foreperiod predictably coupled with removal of the visual fixation stimulus (Saslow 1967). In this gap paradigm, many saccades are initiated with latencies of ~80 ms in macaques and ~100 ms in humans (Boch and Fischer 1986; Boch et al. 1984; Fischer and Boch 1983; Schiller et al. 2004). Because these latencies approach the lower limits imposed by conduction delays in the oculomotor system, these are known as express saccades.

The saccade countermanding task has been widely used to investigate response inhibition (Cabel et al. 2000; Colonius et al. 2001; Godlove and Schall 2016; Hanes and Carpenter 1999; Hanes and Schall 1995; Kornylo et al. 2003; Morein-Zamir and Kingstone 2006; Thakkar et al. 2011; Thakkar et al. 2015; Walton and Gandhi 2006; Wattiez et al. 2016). In this task, subjects fixate a central spot and following a variable amount of time make a saccade to a target stimulus at one of two fixed, spatially separated locations (no-stop trials) presented simultaneously with the disappearance of the fixation dot. On a minority of trials, the fixation point reappeared instructing the monkeys to cancel their planned saccade to the peripheral target (stop-signal trials). Even though express saccades have been reported under overlap conditions (Amatya et al. 2011; Boch and Fischer 1986; Knox and Wolohan 2015), multiple aspects of the stop signal task should discourage production of express saccades. First, because successful performance depends on balancing inhibition and initiation, saccade latencies are typically slower than those observed in other response tasks (Verbruggen and Logan 2009). Indeed, saccade latency increases with the fraction of stop signal trials (Emeric et al. 2007). Second, whereas express saccade production is facilitated by consistent spatiotemporal target presentation, target location and timing are randomized in this task design. Finally, express saccades are most common in gap task condition that encourages release of fixation. In contrast, successful saccade countermanding performance encourages stricter control over visual fixation.

Here we report a serendipitous observation of frequent express saccades produced by monkeys performing a saccade countermanding task. To explore why monkeys produce express saccades in this task, we simulated performance according to the Logan and Cowan (1984) race model. We found that producing a fraction of express saccades can increase reward rate. To verify that express saccade production was motivated by reward contingences, reward for producing express saccades was eliminated. When a minimum saccade latency was enforced for reward, monkeys stopped producing express saccades. We also quantified how saccade latency and express saccade production was affected by the temporal predictability of target presentation. When target presentation could be anticipated, regular but not express saccade latencies decreased. When monkeys performed the countermanding task consistent with the assumptions of the Logan and Cowan (1984) race model, they produced more express saccades than when they performed the task in violation of the assumption. Being incidental findings, their interpretation must be cautious, but they indicate differences in the mechanisms responsible for regular and express saccades, suggest informative future experimental designs, and highlight the range of operation of cognitive control.

## METHODOLOGY

### Animal Care

Data were collected from 3 male bonnet macaques (Macaca radiata, 6.9 to 8.8 kg), and 1 female rhesus macaque (Macaca mulatta, 6.0 kg). Animal care exceeded policies set forth by the USDA and Public Health Service Policy on Humane Care and Use of Laboratory Animals and all procedures were carried out with supervision and approval from the Vanderbilt Institutional Animal Care and Use Committee. Titanium headposts were surgically implanted to facilitate head restraint during eye tracking. Surgical methods have previously been described in detail (Godlove et al. 2011).

### Data Acquisition

Experiments were carried out in darkened, sound attenuated rooms. During testing, monkeys were seated comfortably 43 to 47 cm from a CRT monitor (~48 × 38°, 70Hz) in enclosed polycarbonate and stainless-steel primate chairs and head restrained using surgically implanted head posts. Stimulus presentation, task contingencies related to eye position, and delivery of liquid reinforcement were all under computer control in hard real time (TEMPO, Reflective Computing, Olympia, WA). With the exception of the 70 Hz screen refresh rate, task timing was controlled at 500 Hz. Stimulus sizes and eccentricities were calculated automatically by the stimulus presentation program based on subject to screen distance to allow for increased precision between primate chairs and recording room setups. Stimuli were presented using computer-controlled raster graphics (TEMPO Videosync 640 × 400 pixel resolution, Reflective Computing, Olympia, WA). Stimuli had a luminance of 3.56 cd/m^2^ (fixation point and stop-signals) or 2 cd/m^2^ (targets) on a 0.01 cd/m^2^ background.

### Behavioral Task

Behavior and electrophysiological signals were recorded during the countermanding (i.e., stop-signal) task (Figure 1). Additional details about the behavioral training regime and task have been described previously (Hanes et al. 1998; Hanes and Schall 1995). Data from monkey F was recorded from the first countermanding session. Data from monkey H was recorded after he was well trained on the task. After observations made in an initial 110 sessions, 31 additional sessions were recorded from monkey F to compare the effect of reward contingencies on express saccades. In these sessions, saccade latencies were monitored online. In the first 13 sessions, correct express saccades (i.e. express saccades on no-stop trials) were rewarded. In the following 18 sessions, reward for express saccades was eliminated. Results from this dataset were compared against data recorded several years later from monkey’s Eu and X. Data from both of these monkeys were collected after both were well trained on the task, and neither monkey was rewarded for an express saccade at any point in their training.

**Figure 1:**
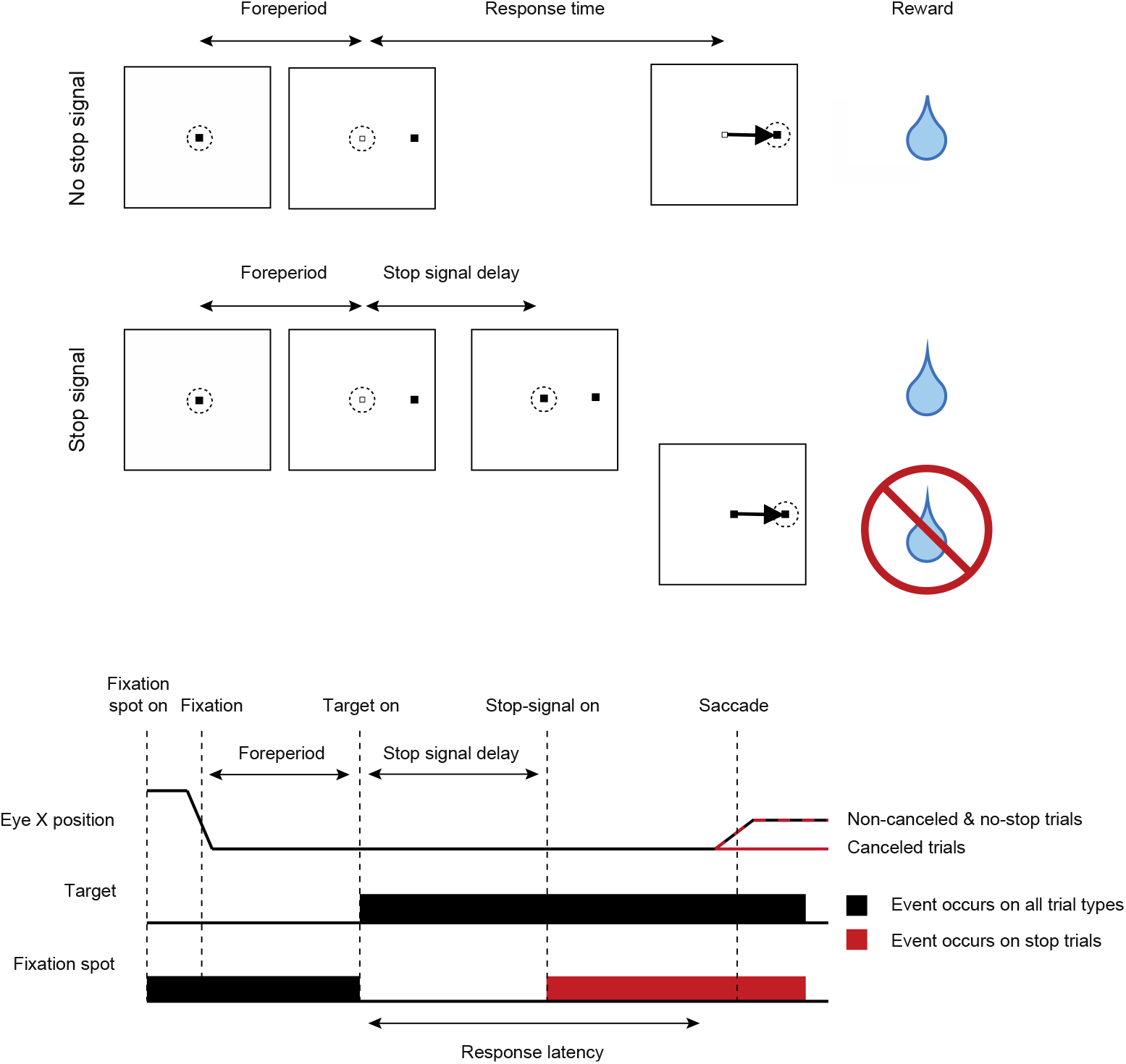
Saccade countermanding task. Monkeys initiated trials by fixating on a central point. After a variable time, the center of the fixation point was extinguished. A peripheral target was presented simultaneously at one of two possible locations. On no-stop-signal trials, monkeys were required to shift gaze to the target, whereupon after a variable period of time, fluid reward was provided. On stop-signal trials (~40% of trials), after the target appeared, the center of the fixation point was re-illuminated after a variable stop-signal delay, which instructed the monkey to cancel the saccade. After holding fixation for a period of time, the monkey received feedback and reward. In a staircase procedure, stop-signal delay was adjusted such that monkeys successfully canceled the saccade in ~50% of trials. In the remaining trials, monkeys made non-canceled errors, in which no reward was delivered. In non-staircased trials, stop-signal delays were randomly selected from a pre-determined set of stop-signal delays.

Trials were initiated when monkeys fixated a centrally presented square which subtended 0.34° of visual angle. After a variable foreperiod, the center of the central fixation point was extinguished leaving a white outline. A target subtending 3° of visual angle simultaneously appeared at 10° to the left or right of the fixation. For two monkeys (F & H) foreperiods were randomly sampled from a uniform distribution. For two other monkeys (Eu & X) foreperiods were randomly sampled from an approximately non-aging, exponentially decaying distribution. On no-stop trials (Figure 1, top), no further visual stimuli were presented. Monkeys were required to make a saccade to the target and hold fixation to obtain reward. Correct trials were rewarded with several drops of juice. On a proportion of trials, the center of the fixation point was re-illuminated after a variable delay providing a “stop-signal” which instructed the monkeys to cancel their impending eye movements and maintain central fixation (Figure 1, bottom). The average proportion of stop trials varied across monkeys for incidental reasons: monkey F: 37%, monkey H: 40%, monkey Eu: 52%, and monkey X: 50%.

On stop signal trials, two trial outcomes were then possible. If monkeys successfully withheld the eye movement and maintained fixation for a period of time (typically 500 ms for monkey F & H, and 1,500 ms for monkey Eu and X), they obtained fluid reward. These trials were designated as “canceled”. If monkeys failed to inhibit the movement, no reward was given, and the trial was termed “non-canceled”. If a saccade was initiated before the stop signal was scheduled to appear, the trial was classified as non-canceled based on the logic of the Logan race model (Logan and Cowan, 1984). No time outs were imposed for a non-canceled error and reward volume was consistent across trial types.

The stop-signal delay (SSD) or time between target and stop-signal presentation determines the probability with which movements can be successfully countermanded (Logan and Cowan 1984). An initial set of SSDs was selected for each recording session based on the experimenter’s knowledge of the animal’s past performance. Although varied from session to session, these SSD’s typically ranged from 43 up to 443 ms, in steps of 16 ms. SSD was then manipulated using either an adaptive staircasing algorithm which adjusted stopping difficulty based on accuracy, or by randomly selecting one of the defined SSD’s. In the staircasing design, when subjects failed to inhibit responses, the SSD was decreased by a random step of 1, 2, or 3 stop-signals increasing the likelihood of success on the next stop trial. Similarly, when subjects were successful in inhibiting the eye movement, the next SSD was increased by a random step of 1, 2, or 3 decreasing the future probability of success. This procedure was used to ensure that subjects failed to inhibit action on ~50% of stop trials overall. Plots showing the probability of responding at each SSD (inhibition functions) were constructed and monitored online to ensure adequate performance.

The median session length for monkey F was 51 minutes (IQR: 27 to 77 minutes), and for monkey H was 236 minutes (119 to 366 minutes). In later data recorded from monkey Eu and X, trial length was fixed. For these monkeys, session lengths ranged from 87 to 184 minutes (modal session time: 169 minutes) for monkey Eu, and from 62 to 193 minutes (modal session time: 120 minutes) for monkey X. For monkey F, the inter-trial interval was fixed (~1000 ms), resulting in a variable trial length. However, for monkey H, the inter-trial interval was varied. In all cases, trial length would be extended if a time-out (~500 ms for monkey F & H, ~5000 ms for monkey Eu and X) was issued if the monkey aborted the trial (i.e. not maintaining fixation on a target, failing to make a saccade before a given deadline). For all sessions, timeouts were not issued for non-canceled trials; instead, the temporal progression of the trial would instead mirror that of a no-stop trial, but with reward omitted.

### Eye Tracking

Eye position data were acquired, calibrated, and streamed to the Plexon computer using the EyeLink 1000 infrared eye-tracking system (SR Research, Ontario, Canada). This system has an advertised resolution of 0.01°. Online, gaze was monitored using digital fixation windows. The size of these windows was determined by the experimenter based on the quality of the eye tracker calibration. Typically, subjects were allowed 1° of stimulus fixation error online. For final trial classification and analysis, saccade initiation and termination were detected offline using a custom algorithm implemented in the MATLAB programming environment (MathWorks, Natick, MA). Saccade starting and ending times were defined as periods when instantaneous velocity was elevated above 30°s^−1^. The eye tracking procedures reliably detected saccades > 0.2° in amplitude. Express saccades were classified as saccades to a target with a latency less than or equal to than 100 ms. No monkey produced enough anticipatory saccades to confound any of the analyses.

## RESULTS

### Monkeys produce express saccades in a countermanding task

We retrospectively examined 425 sessions of saccade countermanding obtained from two monkeys (Monkey F: 110 sessions; Monkey H: 315 sessions). Collectively these monkeys completed 264,422 trials (Monkey F: 77,438; Monkey H: 186,984). While performing this task, monkeys F and H produced saccades with very short latencies which led to bimodal or multi-modal latency distributions. Evident in both single behavioral sessions and across sessions, saccades with latencies ≤ 100ms comprised a separate mode in the saccade latency distributions. These observations are congruent with previous descriptions of express saccades in macaques and were prominent in both monkeys.

Average saccade latencies across all datasets, split by trial type, are displayed in Table 1. On average express saccades comprised 6% of the saccade latencies in a given session, ranging between 0% and 66% of the saccade latencies within a given session. Across 110 sessions, monkey F generated saccades in 38,982 trials (32,992 no-stop, 5,990 non-canceled). The response distribution was clearly multimodal (Figure 2A, top). Median saccade latency across these trials was 248 ms (IQR: 157 – 318 ms). Express saccades were elicited in 4,707 (12.0%) saccade trials and were significantly more common in one of the two target directions (78.73% to 21.27%, one sample t-test, t (106) = 18.369, p < 0.001). The median express saccade latency was 87 ms (IQR: 82 ms – 92 ms). Monkey H generated saccades in 90,347 trials (73,750 no-stop, 16,597 non-canceled). Again, the response time distribution was multimodal (Figure 2A, bottom) with a median saccade latency of 328 ms (IQR: 256 – 420 ms). Express saccades were elicited in 4,061 (4.5%) saccade trials and were significantly more common in one direction (44.05% to 55.95%, one sample t-test, t (297) = −3.628, p < 0.001). The median express saccade latency was 92 ms (IQR: 91.93 ms – 96 ms). Findings for both monkeys are consistent when looking at the single session level (Figure 2B).

**Table 1:** Average ± SD saccade latencies across trial types and data sets.

| Foreperiod |  | Ageing |  | Non-ageing |  |
| --- | --- | --- | --- | --- | --- |
| Saccade type | Trial type | Monkey F<br>(n = 84) | Monkey H<br>(n = 157) | Monkey Eu<br>(n = 12) | Monkey X<br>(n = 17) |
| All | No-stop | 230.7 ± 50.8 | 265.0 ± 50.6 | 311.9 ± 14.2 | 265.8 ± 18.9 |
| | Non-canceled | 238.1 ± 52.8 | 272.5 ± 66.5 | 282.3. ± 25.7 | 232.4 ± 12.9 |
| Regular | No-stop | 252.5 ± 45.7 | 297.8 ± 46.8 | 312.6. ± 14.4 | 266.3 ± 19.1 |
| | Non-canceled | 249.9 ± 46.5 | 301.8 ± 53.8 | 283.8. ± 25.6 | 233.3 ± 13.1 |
| Express | No-stop | 72.1 ± 6.7 | 84.0 ± 3.4 | 68.5 ± 2.5 | 85.5 ± 14.6 |
| | Non-canceled | 71.0 ± 5.5 | 84.1 ± 4.1 | 71.2 ± 8.4 | 85.3 ± 13.1 |

**Figure 2:**
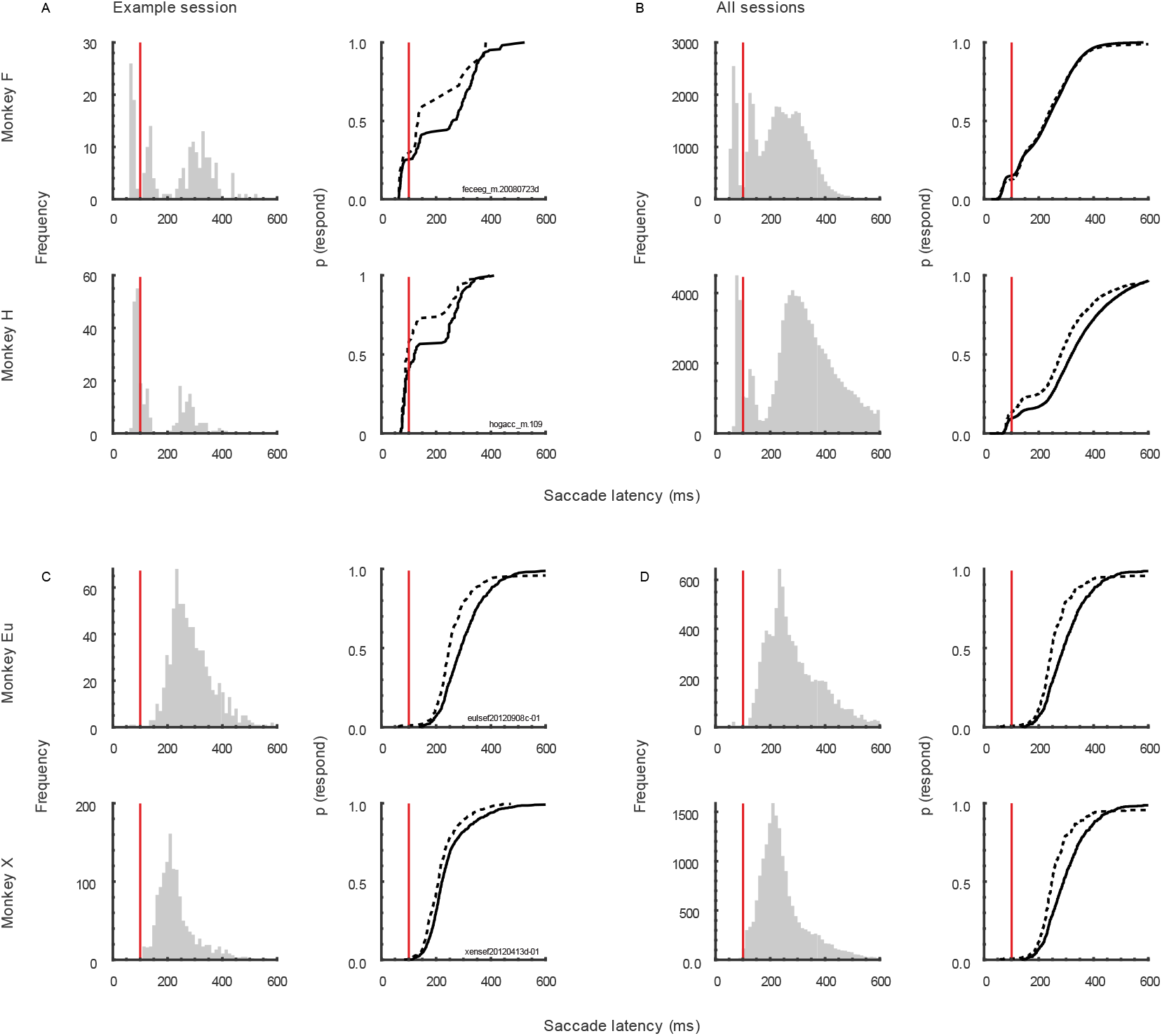
Distributions of saccade latencies during saccade countermanding. Histograms (left) represent saccade latencies on non-canceled and no-stop trials combined, Cumulative distribution functions (right) of saccade latencies are presented for non-canceled (dashed lines) and no-stop trials (solid lines) separately. **A**. Saccade latencies in example sessions for monkey’s F & H. **B**. Saccade latencies collapsed across all sessions for monkey’s F & H. **C**. Saccade latencies in example sessions for monkey’s Eu & X. **D**. Saccade latencies collapsed across all sessions for monkey’s Eu & X. Red vertical lines represent the 100 ms saccade latency mark. Saccades with latencies before this value were classified as express saccades.

### Express saccade production & race model violations

Performance of stop signal countermanding tasks has been explained comprehensively by a race model in which trial outcomes are dictated by the finishing time of stochastically independent GO and STOP processes (Logan and Cowan 1984). The race model is based on the assumption that the finishing time of non-canceled responses (RT _non-canceled_) is consistently faster than the finishing time of responses when there is no stop signal (RT _no stop_). Saccade latencies and express saccade proportions, split by trial types and monkeys, are presented in Table 2, for sessions with and without violations of the race model. In these archived data, we noted a number of sessions in which the RT relationship that justifies the application of the race model was violated. We were surprised to uncover a systematic pattern of variation of express saccade production across sessions when this relationship was or was not violated. We should note that previous publications about countermanding responses based on data from these monkeys utilized sessions in which the race model assumption was respected.

**Table 2:** Average ± SD of saccade latencies and express saccades proportions across trial types and monkeys, split by race model violations.

|  |  | Ageing |  |  |  | Non-ageing |  |  |  |
| --- | --- | --- | --- | --- | --- | --- | --- | --- | --- |
|  |  | Monkey F<br>(n = 84) |  | Monkey H<br>(n = 157) |  | Monkey Eu<br>(n = 12) |  | Monkey X<br>(n = 17) |  |
| RT relationship | Saccade type | RT (ms) | P(Express) | RT (ms) | P(Express) | RT (ms) | P(Express) | RT (ms) | P(Express) |
|  |  | n = 50 sessions |  | n = 83 sessions |  | n = 3 sessions |  | n = 0 sessions |  |
| RT<sub>Non-canceled</sub><br>><br>RT<sub>No stop</sub> | No-stop | 225.8 ± | 0.139 ± | 261.4 ± | 0.151 ± | 303.7 ± | - | - | - |
|  |  | 49.8 | 0.119 | 41.5 | 0.084 | 1.6 |  |  |  |
|  | Non-canceled | 249.8 ± | 0.042 ± | 301.2 ± | 0.110 ± | 316.6 ± | 0.003 ± | - | - |
|  |  | 50.3 | 0.075 | 51.9 | 0.114 | 17.7 | 0.005 |  |  |
|  |  | n = 34 sessions |  | n = 74 sessions |  | n = 9 sessions |  | n = 17 sessions |  |
| RT<sub>Non-canceled</sub><br><<br>RT<sub>No stop</sub> | No-stop | 237.8 ± | 0.105 ± | 267.0 ± | 0.172 ± | 314.7 ± | 0.004 ± | 265.8 ± | 0.003 |
|  |  | 52.2 | 0.138 | 59.3 | 0.100 | 15.5 | 0.004 | 18.9 | (0.003) |
|  | Non-canceled | 221.0 ± | 0.112 ± | 240.3 ± | 0.195 ± | 270.9 ± | 0.009 ± | 232.4 ± | 0.006 |
|  |  | 52.4 | 0.177 | 66.7 | 0.210 | 15.6 | 0.009 | 12.9 | (0.006) |

In these archived data, we observed an unexpected pattern of express saccade production related to satisfying the stochastic independence assumption of the race model. Monkey F violated the RT _non-canceled_< RT _no stop_ relationship in 50/84 sessions (59.5%). On no stop trials, the proportion of express saccades when the relationship was respected was not different from that when the relationship was violated (t (82) = 1.185, p = 0.239, two-tailed). However, during non-canceled trials, the proportion of express saccades when the relationship was respected was significantly greater than that when the relationship was violated (t (82) = −2.495, p = 0.015, two-tailed).

Monkey H showed a similar pattern of behavior and violated the RT _non-canceled_ < RT _no stop_ relationship in 83/157 sessions (52.9%). During no-stop trials, the proportion of express saccades when the relationship was respected was not different from that when the relationship was violated (t (155) = −1.393, p = 0.165). However, on non-canceled trials, the proportion of express saccades when the relationship was respected was significantly greater than that when the relationship was violated (t (155) = −3.223, p = 0.002). Hence, when monkeys F and H performed the countermanding task consistent with the race model assumption of stochastic independence, they produced more express saccades than when they performed the task in violation of the assumption.

Monkey Eu had 3 sessions in which RT _non-canceled_ > RT _no stop_, but the proportion of express saccades did not differ across sessions or trial types. Monkey X had no sessions in which RT _non-canceled_ > RT _no stop_.

### Simulated express saccade production influences behavioral outcomes

Given the apparent inconsistency between performance of the countermanding task and production of express saccades, we sought to understand why monkeys would initiate express saccades. Appreciating that monkeys are motivated to earn fluid reward, we explored whether express saccade production could be advantageous. To do so, we simulated countermanding performance with production of different fractions of express saccades. Unlike previous work (Boucher et al. 2007; Logan et al. 2015), we did not fit parameters based on observed measures of performance. Rather, we simply quantified the amount of reward earned when we simulated countermanding performance while varying the fraction of express saccades produced as well as other parameters of the task. Using this approach, we found increasing express saccade production allowed more trials to be initiated under typical experimental parameters. Furthermore, a small increase in reward rate could be attained by including a small proportion of express saccades.

Saccade latencies were simulated using a linear ballistic accumulator (LBA) to instantiate the independent race underlying countermanding performance (Figure 3A). The saccade GO process was simulated with accumulators having two distributions of rates. One accumulator had slower median accumulation rates, producing longer saccade latencies. The second had faster median accumulator rates, producing express saccades. The fraction of trials governed by the faster accumulator was varied systematically. The STOP process was simulated with another accumulator. On simulated trials with a stop signal, if the STOP accumulator finished before the GO accumulator, then reward was earned for canceling the saccade, and if the STOP accumulator finished after the GO accumulator, then no reward was earned. On simulated trials with no stop-signal, reward was earned for saccades. Model parameters were not fit to performance measures. Instead, we simply explored qualitatively how reward rate varied as a function of various parameters. First, we varied the parameters of GO accumulators to generate different proportions of express saccades (0%, 10%, 50%, 90%; Figure 3B). Second, we varied whether express saccades were rewarded or unrewarded. Third, we varied whether the trial length was fixed or varied. Finally, we varied the proportion of stop signal trials. We examined how timing and experimental parameters similar to those used during data collection influenced average reward and trial rate.

**Figure 3:**
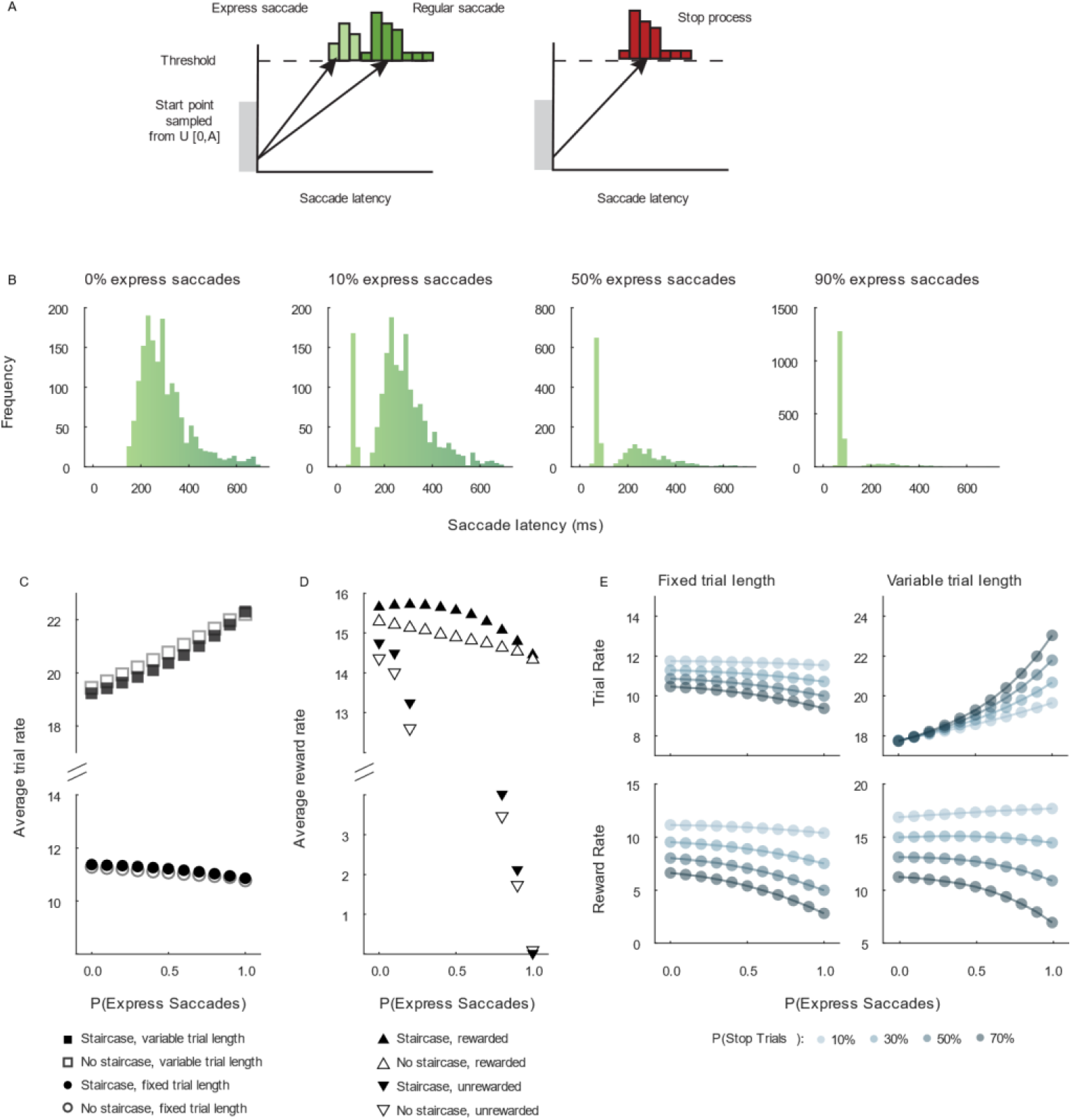
Simulation of reward rate earned with variable fractions of express saccades. **A**. Linear ballistic accumulators for GO process (left) and STOP process (right). The GO process consisted of two mean rates, one producing express saccade latencies, and the other producing regular saccade latencies. Trial outcomes were specified by the finishing time of the fastest process according to the race model (Logan and Cowan, 1984). On trials with no stop-signal, reward was earned after the GO process produced either a regular or an express saccade. On trials with a stop-signal, reward was earned only if the stop process LBA finished before both the regular and the express saccade LBAs. The following combinations of trial parameters were simulated: with or without staircase adjustment of SSD, with variable or fixed trial durations, with or without rewarding express saccade latencies. **B**. Distributions of saccade latencies produced with indicated fractions of express saccades in a simulated session of 1,000 trials. **C**. Average trial length over simulated sessions with (closed) and without (open) staircasing SSD and with variable (square) or fixed (circle) trial lengths. With variable but not fixed trial lengths, trial rate increased with the fraction of… express saccades because more trials could be completed. Whether or not SSD was adjusted in a staircase made negligible difference. **D**. Average reward rate over simulated sessions with (closed) and without (open) staircasing SSD and with express saccades rewarded (upright triangle) or not rewarded (inverted triangle). If express saccades were not rewarded, then reward rate decreased dramatically with increasing fraction of express saccades, whether or not SSD was adjusted in a staircase. If express saccades were rewarded, then reward rates were higher and decreased less with increasing fractions of express saccades. However, if SSD was adjusted in a staircase, then reward rate was maximal when ~10% express saccades were produced. **E.** Variation of trial rate (top) and reward rate (bottom) as a function of proportion of express saccades for fixed (left) and variable (right) trial durations. Progressively darker points plot values with progressively higher fraction of stop-signal trials. With lower fractions of stop-signal trials, the cost of express saccades on reward rate is reduced.

### Express saccade production increases the number of trials available

Under a fixed intertrial interval, trial length can vary as a function of response time. Under this paradigm, a faster saccade latency may lead to a shorter trial length, and thus provide the monkey with more trials per minute in order to gain reward. With 40% stop trials, simulated data demonstrated that an increase in the proportion of express saccades led to an increase in trial rate (number of trials per minute) (Figure 3C, squares). However, when trial lengths are fixed, trial rates only slightly decrease and subjects can only initiate around 11 trials per minute (Figure 3C, circles).

Given that there is no penalty for making express saccades in no-stop trials, but there are time-out penalties in stop-trials, an increase in stop trial proportion will increase the error rate and trial length and reduce overall trial rate. As such, we examined if these values varied as a proportion of stop trials in a given session (Figure 3E, upper panels). Interestingly, if a session had a lower proportion of stop-trials (~10%), then producing more express saccades led to only a small increase in the number of trials available. However, if a session had a greater proportion of stop-trials (~70%), then producing more express saccades led to a greater increase in the number of trials available compared to lower stop-trial proportions. This is only true for variable trial lengths. If trial lengths were fixed, then express saccades decreased trial rates for all stop-signal proportions.

### Express saccade production is not detrimental to reward rate

When trial length varied, express saccades were rewarded, and stop-signal delays adapted in a staircase procedure, we found systematic variation in reward rate with the proportion of express saccades in a session (Figure 3D). At typical stop-trial proportions (~40%), producing a small proportion of express saccades was not detrimental to performance, and led to slightly increased reward rates (Figure 3D). This relationship changed dependent on the proportion of stop-trials in a given session (Figure 3E, lower panels). If a session had a lower proportion of stop-trials (~10%), then producing more express saccades lead to a higher reward rate (Figure 3E). However, if a session had a greater proportion of stop-trials (~60%), then producing more express saccades was detrimental to performance (Figure 3E). This is only true for variable trial lengths. If trial lengths were fixed, then express saccades were detrimental to reward rates for all stop-signal proportions.

### Experimental express saccade production varies with reward contingencies

The findings from the simulation demonstrated express saccade production can influence trial and reward rates under typical experimental conditions and parameters. Given this relationship, we then looked at these features within the data. In this experimental data, express saccade production increased from early to later training sessions and was mirrored by an increase in average reward rates. When express saccades were no longer rewarded, their prevalence significantly decreased. Two other monkeys trained when express saccades were unrewarded produced very few express saccades across 10,000’s of trials.

### Express saccade production is learnt through training

For monkey F, we tracked the progression of express saccade production from the initial training session onwards. We observed the average proportion of express saccades significantly increased over time (r = 0.49, p < 0.001). In parallel, the average reward rate within each session increased significantly with experience (r = 0.34, p < 0.001). This pattern is also clear when sessions were divided into five equal groups. From the earliest to latest training period, the proportion of express saccades increased significantly (F (4,171) = 7.76, p < 0.001, Figure 4A), as did reward rate (F (4,171) = 11.69, p < 0.001, Figure 4B).

**Figure 4:**
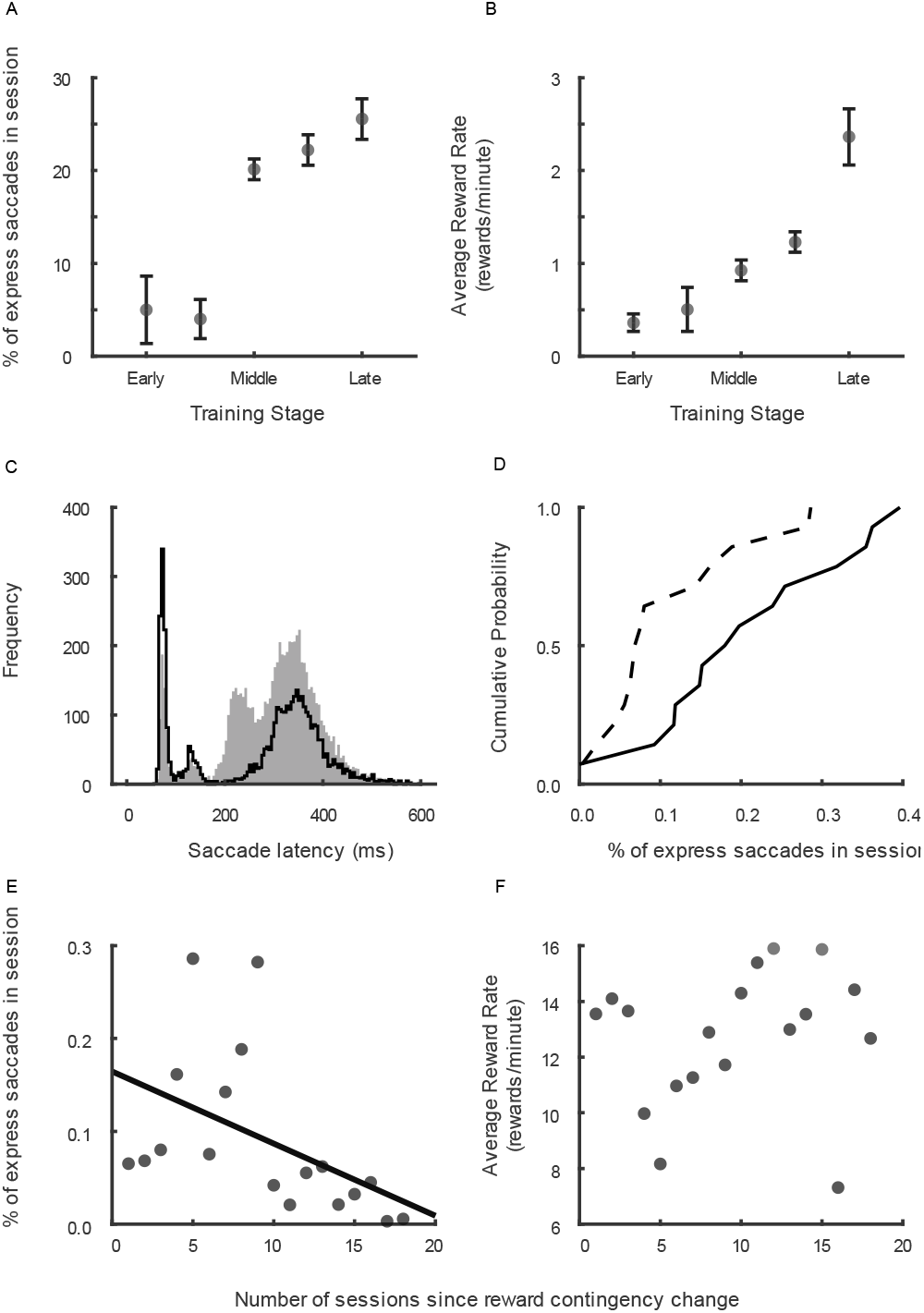
Learning and unlearning express saccades. Variation of performance of monkey F as a function of training and task contingencies. **A**. Average reward rate earned from early to late stages of training in 18 sessions when express saccades were rewarded just like regular saccades on no stop trials. **B**. Percentage of express saccades produced across corresponding stages of training. The monkey exploited contingencies of the task to earn more reward by producing a fraction of express saccades. **C**. Distributions of saccade latencies when express saccades were rewarded (open) and in subsequent sessions when only no stop trials with regular saccade latencies were rewarded (filled). When express saccades were not rewarded, the monkey produced more regular latency saccades to earn reward. **D**. Cumulative probability of express saccade production in sessions when they were rewarded (solid) and in later sessions when they were not (dashed). **E**. The percentage of express saccades decreased across sessions following changing contingency to not reward express saccades. **F.** Average reward rate did not decrease across corresponding sessions following the change of reward contingency.

### Express saccade production is reduced when no longer rewarded

To causally test the association between reward rate and express saccade production, we recorded 31 additional behavioral sessions with monkey F. These sessions were recorded after the 425 sessions previously described and were not included in the previous analyses. In these sessions, we monitored saccade latencies online. We first recorded a block of sessions where correct express saccades were rewarded (n = 13 sessions). This was followed by a block of sessions in which express saccades were unrewarded (n = 18 sessions).

Sessions with both unrewarded and rewarded express saccades had multimodal saccade latency distributions (Figure 4C). When rewarded, express saccade trials comprised 22.6% of the latency distribution. However, when unrewarded, express saccades became much less common and a new mode appeared in the saccade latency distribution in the 200 – 250 ms range. In these sessions, the proportion of express saccades dropped considerably and comprised only 7.9% of the latency distribution, χ^2^ (1, N = 10465) = 405.648, p < 0.001 (Figure 4D). Reverse to observations in the early training stages (when express saccades were rewarded), we found that the average proportion of express saccades significantly decreased with time from the reward manipulation (r = −0.723, p < 0.001, Figure 4E). Interestingly, average reward rate exhibited no trends (r = 0.183, p = 0.467, Figure 4F).

### Monkeys trained with no reward contingencies don’t produce express saccades

Further evidence that monkeys learn to produce express saccades through training was obtained by examining the saccade latency of two other monkeys who were trained with unrewarded express saccades. We examined an additional 29 sessions of saccade countermanding in two other monkeys (Monkey Eu: 12 sessions; Monkey X: 17 sessions) to compare training histories and reward contingencies. Collectively, these monkeys completed 33,816 trials (Monkey Eu: 11,583; Monkey X: 22,233). The saccade latency distributions were unimodal for both monkeys (Figure 2C & D, for all sessions and an example session respectively). Express saccades were elicited in only 107 trials, comprising only 0.40% of all saccade latencies. Across the 29 sessions, monkey Eu produced 42 (0.49%) express saccades and monkey X produced 65 (0.36%) express saccades.

### Temporal predictability can affect express saccade production

Express saccades occur more often when the timing of a target presentation is predictable (Pare and Munoz 1996; Saslow 1967; Schiller et al. 2004). As such, we examined saccade latency as a function of the foreperiod between fixation at the central cue and target presentation. We started by looking at the distributions and respective survivor functions for foreperiods in our study. The survivor function is just the probability that a target has not yet appeared by a given time. When foreperiods are sampled from a uniform distribution, the survivor function linearly decreases. Hence, as time passes, the proportion of the distribution from which the target onset time can be selected decreases. Although this function offers some insight to the temporal evolution of the distribution, temporal predictability is quantified by the hazard function, which is the conditional probability of an event occurring at a given time given that it has not yet occurred (Luce 1986; Nobre et al. 2007). Formally, the hazard rate is the ratio of the probability density of the event divided by the survivor function. In uniform distributions, the hazard rate for target presentation increases over time, resulting in an ageing function. This ageing function stipulates target appearance becomes predictable as the foreperiod progresses. Conversely, when sampled from exponentially decaying distributions, the hazard rate is invariant over time. Under these conditions, the time of target onset is unpredictable. Given the retrospective nature of this study, we first examined the various patterns of foreperiod distributions experienced by monkeys within our dataset. Although the pattern of foreperiods across some sessions for monkey F and H were variable, we identified a subset of sessions for each monkey with a reliable, uniform foreperiod distribution. Results for regular and express saccades separately are highlighted in Table 3.

**Table 3:**
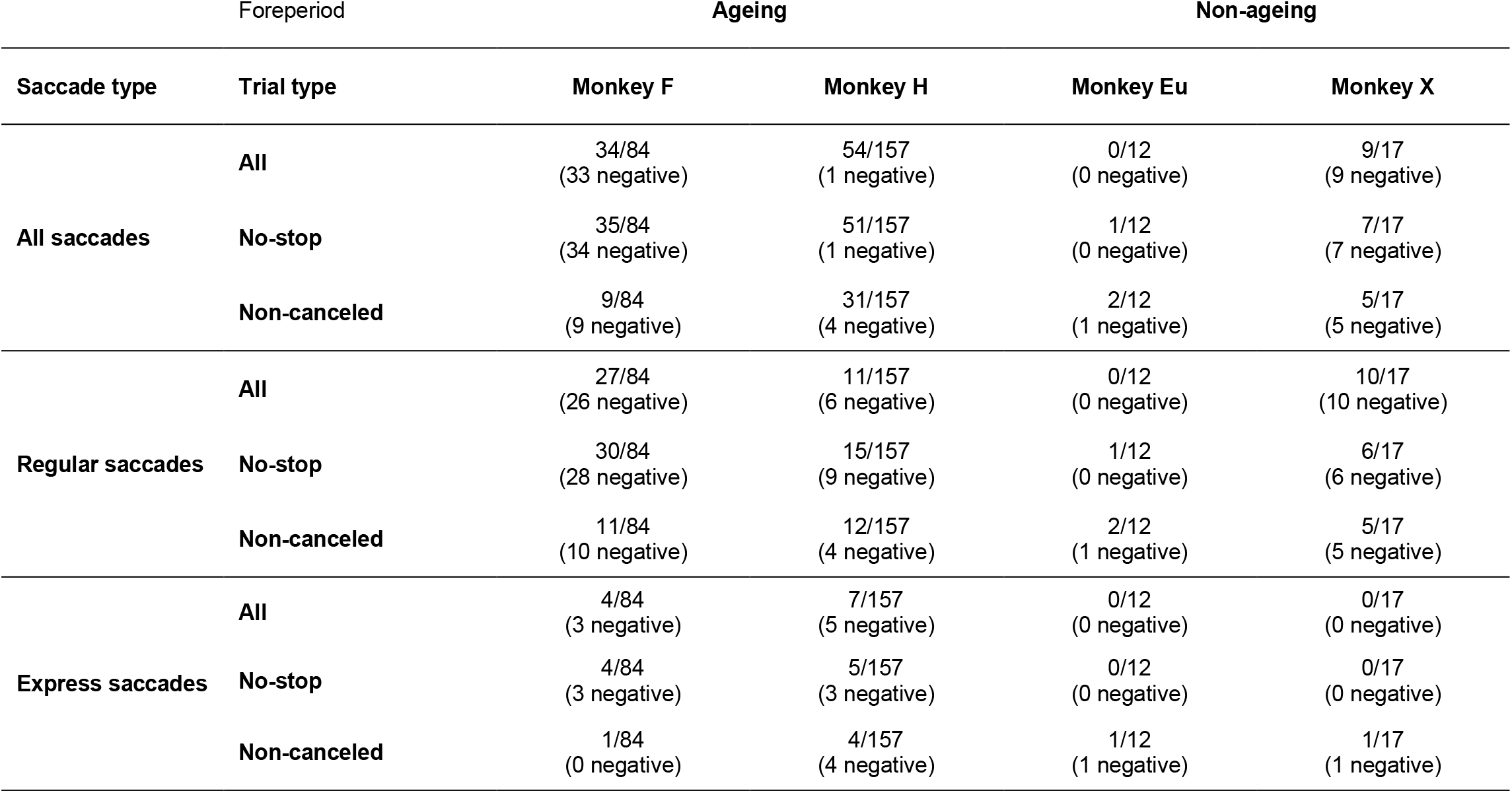
Number of significant correlations between saccade latency and foreperiod duration for each foreperiod distribution, each saccade type (all, regular, or express), and each trial type (all, no-stop, and non-canceled).

In the majority of sessions for monkey F (84/110, 76%) a uniform distribution of foreperiods ranged from ~420 to ~630ms (Figure 5A). A linear decrease in the survivor function is associated with a continually increasing hazard rate of times when the foreperiod elapsed. In some of these sessions, saccade latency decreased with foreperiod duration (33/84 sessions, 39%). However, despite the decrease of saccade latencies at longer foreperiods, the proportion of express saccades did not vary over foreperiod duration (F (2, 249) = 2.11, p = 0.123).

**Figure 5:**
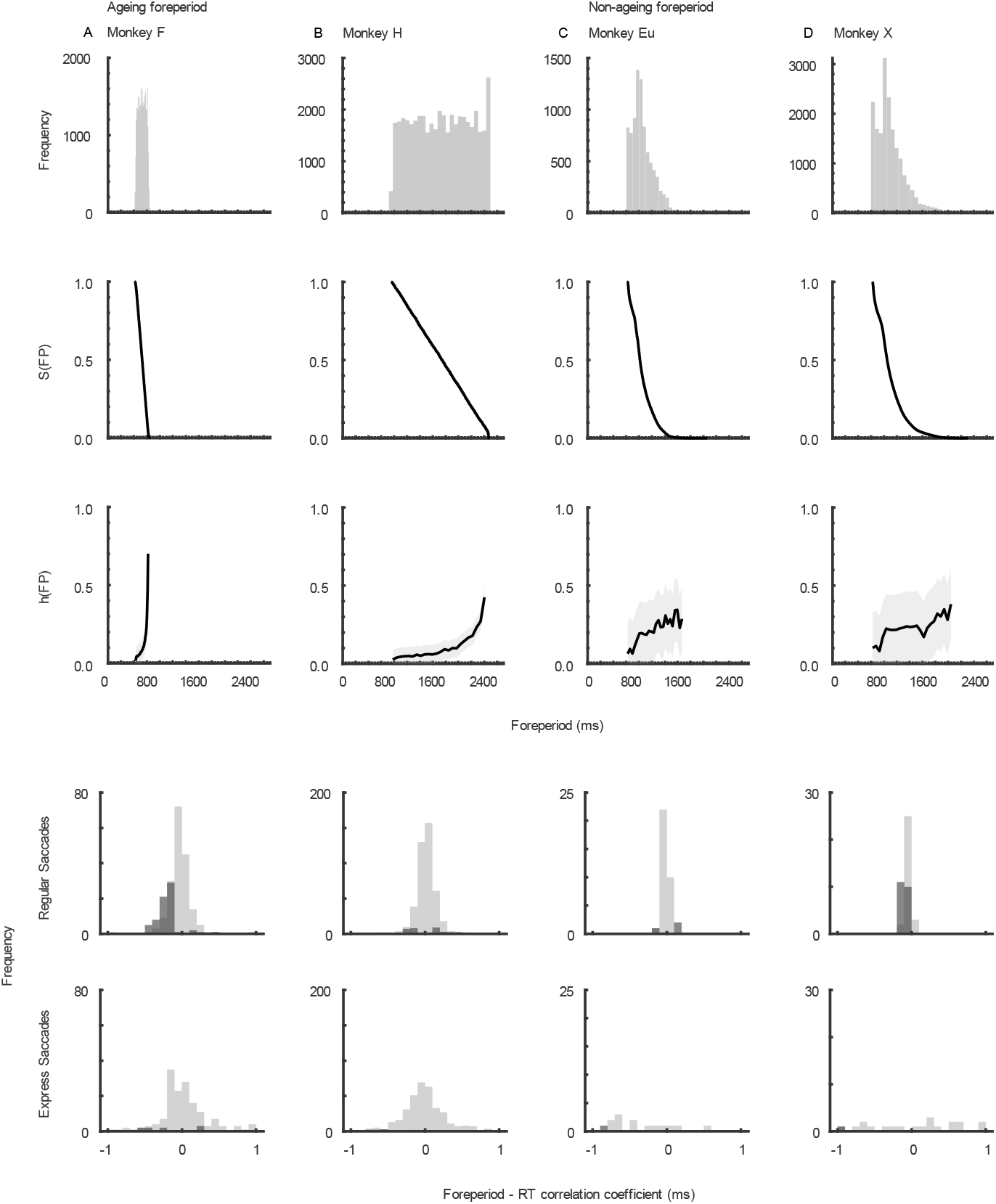
Distributions of foreperiods and relation to saccade latency. Histograms (*1^st^ row*), survivor distribution (*2^nd^ row*), and hazard rate (*3^rd^ row*) of foreperiods used for monkey F (**A**), monkey H (**B**), monkey Eu (**D**), and monkey X (**E**). Monkey’s F and H experienced aging foreperiods, and monkey’s Eu and X experienced approximately non-aging foreperiods. Correlation coefficients for the correlation between saccade latency and foreperiod are presented in the *bottom row*. Dark shades represent sessions in which the correlation coefficients were significant; light shades are non-significant sessions. Significant foreperiod effects were limited to regular saccades for monkey F and Eu. Foreperiod duration did not influence express saccade latency.

We observed the opposite pattern of results from monkey H, who experienced a uniform distribution of foreperiods in half of the sessions (157/315 sessions, 50%) ranging over a much wider interval, ~760 to ~2275 ms (Figure 5B). No relation between saccade latency and foreperiod duration was found in most of these sessions (104/157, 66%), but in the remainder monkey H demonstrated increasing latencies with longer foreperiods (53/157, 34%). This slowing of saccade latencies was accompanied by reduced express saccades production (F (15, 2438) = 14.44, p < 10^−5^).

Monkey Eu and monkey X experienced a common pattern of foreperiod distribution ranging from ~600 to ~ 2100 ms (n = 29/29 sessions). Compared to the foreperiod experienced by monkeys F & H, the survivor function of this distribution was exponentially decaying, resulting in a non-ageing foreperiod (Figure 5C & D). As expected with non-ageing foreperiods, none of monkey Eu’s sessions had significant changes in saccade latency or in the proportion of express saccades generated as a function of foreperiod. However, and somewhat unexpectedly, a foreperiod effect was observed in over half of the sessions for monkey X (9/17, 53%). Furthermore, although uncommonly produced, the proportion of express saccades did decrease as a function of foreperiod bin, F (15, 206) = 2.26, p = 0.006.

## DISCUSSION

We observed multimodal distributions of saccade latencies in a saccade countermanding task. Furthermore, simulations revealed that trial and reward rate in this task can be increased by varying the proportion of express saccades made within a session, under certain conditions. We found that monkeys learnt and exploited this association over time. We then manipulated the reward contingencies for express saccade production and found when monkeys made significantly lower proportion of express saccades when no longer rewarded for them. It is important to note the incidental nature of these findings, and that the features studied were not the main focus of the investigations. As such, our interpretations of these data are limited and future experimental designs should be employed to test them more thoroughly. However, this study has demonstrated for the first time how express saccade production can be accomplished through strategic modifications in saccade latencies, under the guidance of cognitive control.

### Express saccades production in atypical conditions

Consistent with classic reports of express saccades in monkeys, the earliest modes in the resulting distributions peak below 100ms and display very little variance (Boch and Fischer 1986; Boch et al. 1984; Fischer and Boch 1983; Schiller et al. 2004). Although express saccades have also been observed in “overlap” conditions, where the fixation spot remains on even whilst the target is illuminated (Amatya et al. 2011; Boch and Fischer 1986; Knox and Wolohan 2015), this finding in the saccade countermanding task was unexpected given the range of other features that are counterproductive for express saccade production. As such, although intended to be random and unpredictable, the production of express saccades under these conditions may suggest monkeys have some knowledge about the underlying contingencies in our experiment, and generated make predictions to adapt saccade latencies accordingly.

### Temporal predictability and saccade latency

Monitoring the timing of events allows for temporal predictions of target onset to be generated, allowing for a movement to be planned and prepared ahead of time (Kolling and O’Reilly 2018; Petter et al. 2018). The use of hazard functions is motivated by the premise that participants have a representation of the elapsed time in the current situation based on knowledge of the experienced distribution of durations. If an interval elapsing before an event is randomly sampled from a uniform distribution (also known as an aging distribution), then participants will behave as if the event becomes more likely as time progresses. Thus, temporal prediction is critical in reaction time tasks in which foreperiods are a major determinant of response latency. Using distributions of foreperiods selected uniformly from a range of values, i.e., aging, manual response time studies have shown as foreperiod increases, response times become faster (Ameqrane et al. 2014; Correa and Nobre 2008; Drazin 1961; Naatanen 1970; Niemi and Näätänen 1981; Thomaschke et al. 2011). Consistent with the aforementioned manual response studies, this finding has been replicated focusing on saccade latencies in humans (Findlay 1981) and monkeys (Schall 1988). We replicated this effect in the current study. Monkeys can make effective, but idiosyncratic temporal predictions about the time a target may appear on screen.

### Reward maximization

The results demonstrate that monkeys adjusted saccade production to maximize earned rewards. By producing express saccades, monkeys could increase the number of trials available and thus the opportunity to gain a reward. Monkeys did not produce enough express saccades to be detrimental to reward rate.

If the total time of the task is fixed, a rational subject will trade off speed and accuracy in order to try and maximize the amount of reward they can gain in the given session. If more time is spent on one trial, there will be a greater chance of a correct response; however, longer trials reduce the total number of trials available and thus the total amount of reward available. This ultimately gives rise to a speed-accuracy trade-off which aims to maximize gains through flexible cognitive control of behavior.

Non-human primates are adept at optimizing reward rate by exploring different behavioral strategies and task parameters (Feng et al. 2009). Feng et al. (2009) developed a model allowing for the calculation of choice bias that yields the optimal harvesting of reward, given the animals’ sensitivities to a visual stimulus. They found monkeys acquire over 98% of the possible maximum rewards, with shifts away from optimality erring in the direction of smaller penalties. In addition to this, monkeys have also shown to discover unexpected ways to exploit task contingencies. Lowe and Schall (2019) demonstrated this effect in a pro-/anti-saccade visual search task where a vertical singleton cued that a pro-saccade should be made towards it, and a horizontal singleton cued an anti-saccade away from it. Close examination of the reward contingencies of the experiment found that shifting gaze towards a vertical stimulus was the correct outcome on 66% of trials. Exploiting this contingency, they found that one monkey produced more frequent and faster responses to vertical items than to any other item in the array, suggesting they adopted a strategy of searching for vertical items opposed to using the stimulus response rule provided by the singleton. Bichot et al. (1996) found a similar behavior in a color pop-out task: monkeys trained exclusively to find one color in an array persistently direct gaze to stimuli of that color, regardless of whether it is a distractor or target. These are just some of the examples of macaque monkeys creatively exploiting reward contingencies.

### Conclusion

We report incidental findings revealing that express saccade production can be adjusted to increase the opportunities to gain reward. This observation suggests that the mechanisms of express saccade generation are sensitive to trial history and integrate information over multiple trials. This suggests that higher cortical areas that are involved in performance monitoring may contribute to the executive control of express saccade production (Dash et al. 2020; Donahue et al. 2013; Sajad et al. 2019; So and Stuphorn 2010; Stuphorn and Schall 2006; Stuphorn et al. 2000). Further research is needed to understand whether express saccade production can be controlled independently or whether express saccade production is accomplished as a corollary to strategic adjustments in overall RT.

## ACKNOWLEDGEMENTS & FUNDING

The authors thank K. A. Lowe and J. A. Westerberg, and Dr.’s T. Reppert, A. Sajad, and C.R. Subraveti, for helpful discussions and comments on the manuscript. This work was supported by R01-EY019882 (SE), R01-MH55806 (JS), P30-EY08126 (JS), and by Robin and Richard Patton through the E. Bronson Ingram Chair in Neuroscience (JS). There are no reported conflicts of interest.

